# NMDAR-dependent emergence of behavioral representation in primary visual cortex

**DOI:** 10.1101/688366

**Authors:** Alicja Puścian, Hadas Benisty, Michael J. Higley

**Affiliations:** Department of Neuroscience, Kavli Institute of Neuroscience, Yale University School of Medicine, New Haven, CT 06510 USA

## Abstract

Neocortical sensory areas are generally thought to faithfully represent external stimuli. However, a growing body of evidence suggests that cortical networks exhibit considerable functional plasticity over multiple temporal scales, allowing them to modify their output to reflect ongoing behavioral demands. Nevertheless, the dynamics of sensory and non-sensory representations during acquisition of stimulus-guided behavior are not well understood. We performed longitudinal 2-photon imaging of activity in primary visual cortex (V1) of mice learning a conditioned eyeblink task. We found that both excitatory and inhibitory neurons robustly encode the visual stimulus throughout training despite a significant experience-dependent reduction in response magnitude. In contrast, both pyramidal neurons and parvalbumin-expressing interneurons exhibit emergence of behavioral representation during learning. The plasticity of visual response magnitude and behavioral representation is abolished following loss of NMDA-type glutamate receptors. Overall, our findings demonstrate that V1 networks can dynamically multiplex distinct behaviorally relevant representations over the course of learning.

## Introduction

Primary sensory areas of the mammalian neocortex, including the primary visual cortex (V1) have traditionally been thought to faithfully represent features of external stimuli, with minimal influence of intrinsic factors such as behavioral state or motivation. However, a wide literature has now established that such representations can be highly plastic over a range of temporal scales. For example, moment-to-moment changes in arousal level produce robust modulation of neuronal firing in V1 and alterations in perceptual ability ^1–4^. Additionally, learning associations between sensory stimuli and behaviorally relevant outcomes can drive changes in neuronal structure, activity patterns, and perceptual ability ^5–14^.

Most previous investigations of functional plasticity in V1 during learning have focused on modifications of sensory representations by neuronal activity. In both primates and rodents, repeated association between a visual stimulus and reward results in modification of feature selectivity (e.g., orientation preference) by single neurons ^6–8, 10–15^. Such representations are also strongly shaped by top-down influences reflecting levels of arousal or attention ^3, 4, 16^, and prior stimulus expectation ^17, 18^. Beyond encoding visual stimulus features, emergence of neuronal activity corresponding directly to behavioral choice or trial outcome has also been described ^6, 19–21^. Similar results have also been observed in other sensory modalities ^22–25^. Overall, these findings suggest that a strong convergence of behaviorally relevant signals occurs, even at the earliest stages of cortical sensory processing. Nevertheless, the dynamic representation of both sensory and non-sensory information across learning by varied populations of excitatory and inhibitory neurons has not been well-characterized. Moreover, the cellular mechanisms driving such functional plasticity are largely unknown.

Several groups have shown that classical eyeblink conditioning provides an excellent model for investigating the neural correlates of sensorimotor learning ^26–29^. We recently showed that mice rapidly learn to form associations between visual stimuli and aversive corneal air-puffs, resulting in expression of a conditioned blink response ^21^. Neuronal activity within V1 is required for task performance and significantly predicts behavioral outcome in expert animals ^21^. Here, we applied chronic 2-photon imaging of targeted neuronal populations within mouse V1 to investigate the dynamics and plasticity of sensory and behavioral representations throughout learning. Our results demonstrate that visual experience drives a strong reduction in sensory-evoked activity while preserving the ability of excitatory and inhibitory cells to robustly encode the presence of a sensory stimulus. In contrast, both single neurons and the overall population exhibit significant emergence of behavioral representations over the same period. Activity of pyramidal neurons (PNs) and parvalbumin-expressing interneurons (PV-INs), but not somatostatin- or vasoactive intestinal peptide-expressing interneurons (SOM-INs, VIP-INs), is predictive of behavioral performance on individual trials. Finally, we show that the plasticity of both visually evoked responses and behavioral representation requires expression of NMDA-type glutamate receptors (NMDARs), suggesting a critical role for synaptic plasticity in the emergence of task-relevant activity in V1.

## Results

To study the relationship between V1 activity and the acquisition of sensory-guided behavior, we developed a visually cued eyeblink conditioning task. Briefly, head-fixed mice placed on a freely-moving wheel learned to associate a contrast-modulated drifting grating (conditioned stimulus, CS) with an ipsilateral corneal air-puff (unconditioned stimulus, US) that elicits an unconditioned blink response (UR). Repeated stimulus pairings result in a conditioned blink response (CR) that occurs after the onset of the visual stimulus but prior to the air-puff (Figure 1a, see Online Methods). Video recording of the eyelid allowed us to easily determine correct (presence of a CR) versus incorrect (presence of a UR only) performance on individual trials. Mice (n=7) readily learned this task over a few days with a consistently low false-alarm rate (spontaneous blinks, Figure 1b). Acquisition of this behavior did not alter the magnitude or latency of the CR (Figure 1c). Importantly, we previously showed that learning and performance of this task requires intact V1 circuits ^21^. For subsequent analyses of neural imaging data, we divided the 2-week training period into three phases (early, mid, and late) based on average performance (Figure 1b-c).

**Figure 1.**
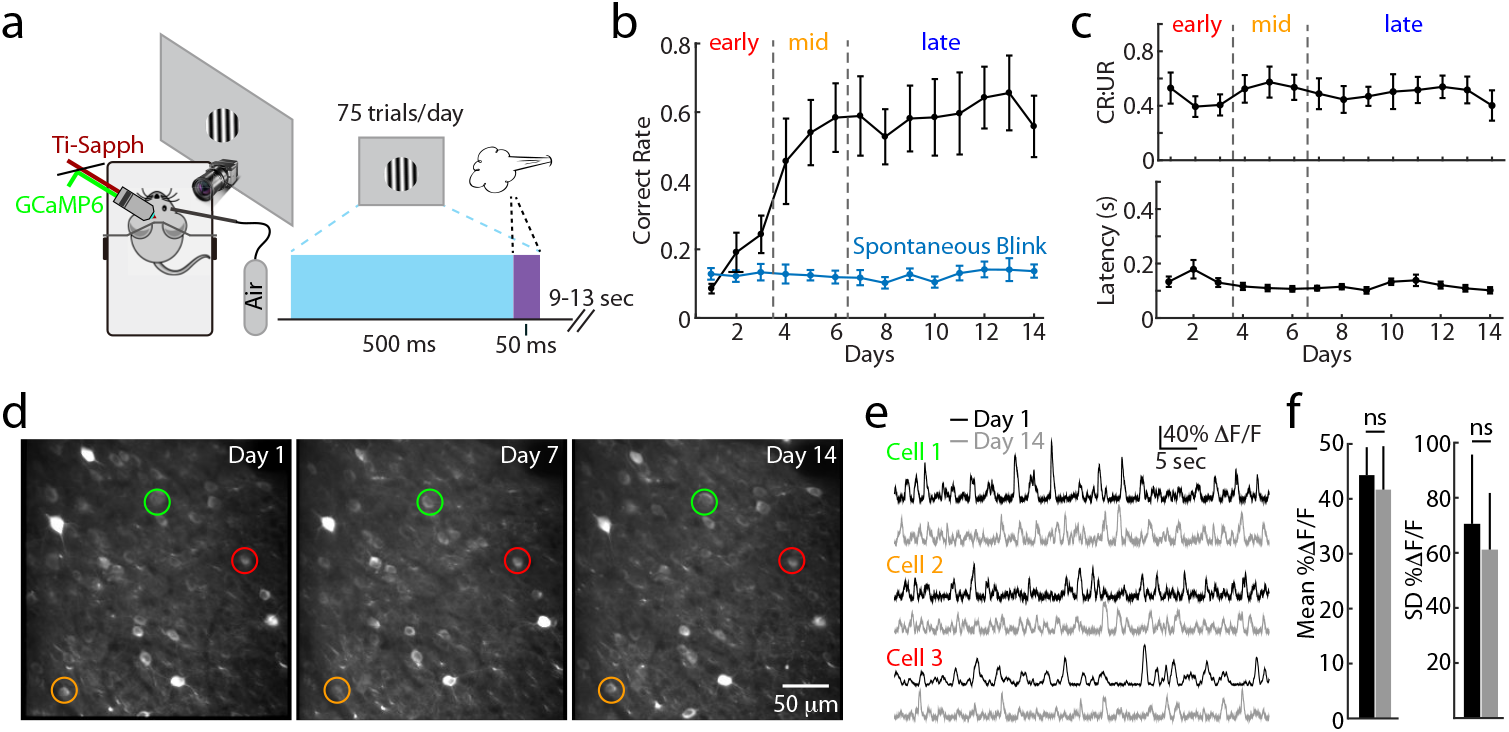
Longitudinal imaging of neuronal activity during visually cued eyeblink conditioning. **a**, Schematic illustration of the behavioral setup (left) and trial structure (right). **b**, Performance over learning for the layer 2/3 PN imaging cohort (n=7 mice, black). Dots with error bars represent average ± SEM. Spontaneous blink rate per 450 ms period is shown (blue). Training phases were divided into tertiles (early, mid, and late) based on average group performance. **c**, Magnitude of the conditioned blink response normalized by the unconditioned response (CR:UR) and blink latency over learning. Dots with error bars represent average ± SEM (n=7 mice). **d**, Example in vivo 2-photon imaging of layer 2/3 neurons expressing GCaMP6s, collected on three different training days from the same mouse. **e**, Example traces showing spontaneous activity from the three color-coded cells indicated in (d) for Day 1 (black) and Day 14 (gray) of training. **f**, Mean (left) and standard deviation (right) of spontaneous activity (%ΔF/F) averaged across all layer 2/3 neurons, for Day 1 (black) and Day 14 (gray). Bars represent average ± SEM (n=7 mice). ns indicates p>0.05, paired t-test.

We then investigated the dynamics of V1 activity during task acquisition. We used a neuron-specific AAV vector to drive expression of the genetically encoded calcium indicator GCaMP6s in layer 2/3, where the large majority of cells are excitatory pyramidal neurons (PNs). We then imaged the activity of putative PNs during learning via 2-photon microscopy. This approach enabled us to reliably track the same neurons longitudinally across each day of the 2-week training period (Figure 1d). Neuropil-subtracted fluorescence signals (ΔF/F) for each neuron in the field of view were collected and analyzed over time (see Online Methods). Repetitive imaging produced no significant change in the mean (44.3±5.1% vs. 41.7±7.9%, n=7 mice, paired t-test, p=0.797) or standard deviation (70.5±25.2% vs. 61.2±20.6%, n=7 mice, paired t-test, p=0.817) of spontaneous fluorescence signals for individual cells over two weeks (Figure 1e-f), suggesting this protocol did not significantly perturb baseline neuronal excitability or negatively impact cell health.

We analyzed visually evoked responses throughout the 2-week training period (Day 1-Day 14). In order to make comparisons with various control cohorts (see below), we also added an additional day before (Day 0) and after (Day 15) training in which the visual stimulus was presented alone (Figure 2a). As we showed previously ^21^, spontaneous blinks can evoke responses in V1 neurons (Supplemental Figure 1a). To exclude any potential contamination of measurements of stimulus-evoked responses by this activity, we restricted our analyses to a 300 ms period following stimulus onset (Supplemental Figure 1b). Within this brief temporal window, we found the average %ΔF/F to be significantly more variable than the calculated linear slope (%ΔF/sec) across trials, reflecting sensitivity to spontaneous signal fluctuations. However, the mean values for these two parameters were highly correlated, suggesting they reflect the same underlying activity (Supplemental Figure 1c-d). For this reason, we use slope as the measure of response magnitude in subsequent analyses.

**Figure 2.**
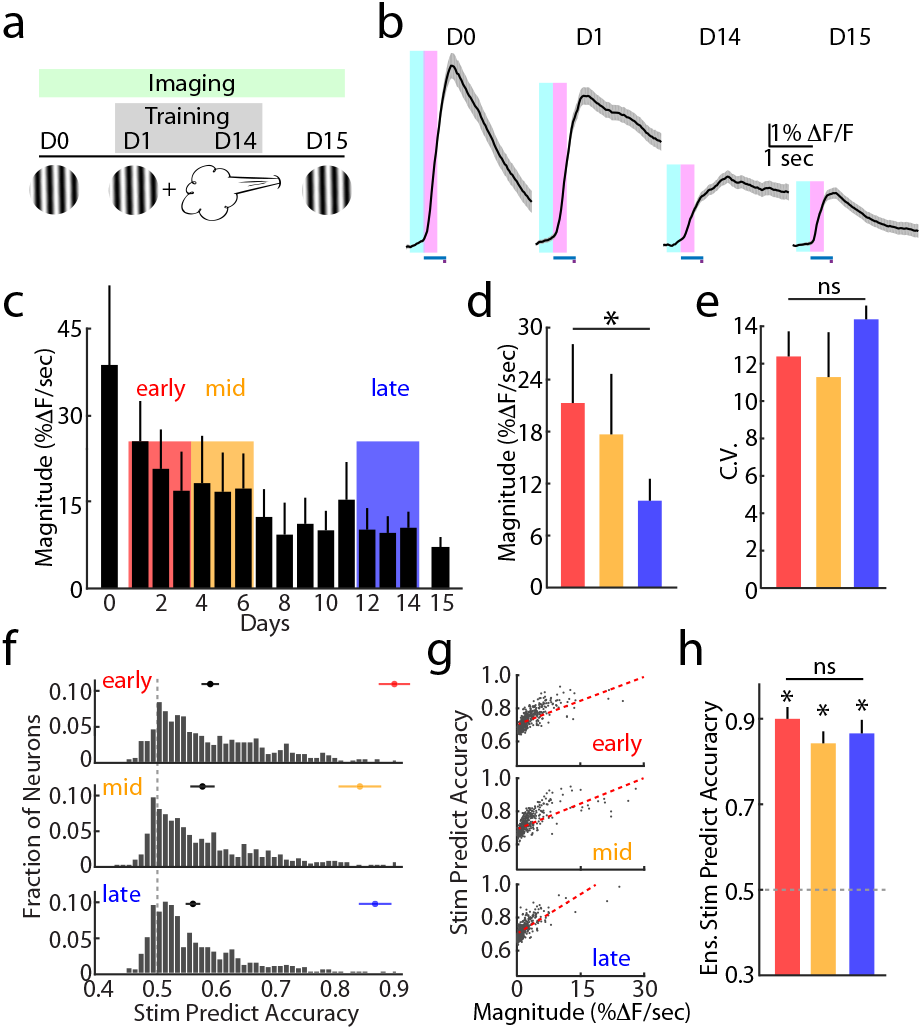
Training reduces visual response magnitude without altering stimulus predictability. **a**, Schematic illustration of the stimulus presented for each day of training (CS alone on Day 0 and Day 15, CS+US on Days 1-14). **b**, Average visually-evoked response for one example layer 2/3 neuron across four different training days. Timing of visual stimulus (blue bar) and air-puff (purple bar) are shown below each trace. Intervals for measuring baseline activity (light blue window) and visual response magnitude (pink window) are shown superimposed. Lines and shading indicate average ± SEM across all trials for the given day. **c**, Population values for the visual response magnitude (given as %ΔF/sec) for each day of training. Analysis windows (3 days) for early (red), mid (orange), and late (blue) training phases are indicated. **d**, Average population values for response magnitude within each training phase, corresponding to colors in (C). Bars represent average ± SEM (n=7 mice). * indicates p<0.05, paired t-test for early vs. late. **e**, Average population values for the coefficient of variation for visual responses within each training phase. Colors as in (c). Bars represent average ± SEM (n=7 mice). ns indicates p>0.05, paired t-test for early vs. late. **f**, Distribution of stimulus prediction accuracy values using a linear decoder for responses of individual layer 2/3 neurons across each training phase. Chance level (0.5) is indicated (gray dashed line). Black circles indicate average ± SEM for the population of individual neurons. **g**, Relationship between stimulus prediction accuracy and response magnitude for individual neurons across each training phase. Red dashed line indicates Spearman’s rank correlation. **h**, Average stimulus prediction accuracy values using a linear decoder for the ensemble activity. Colors as in (c). Bars represent average ± SEM (n=7 mice) and are also indicated by colored circles in (F). * indicates p<0.05, t-test relative to chance. ns indicates p>0.05, paired t-test for early vs. late.

Visual eyeblink conditioning was associated with a dramatic reduction in the evoked response over the course of training (Figure 2b-c). This result was significant when comparing response magnitude between early and late learning phases (21.0±6.9% vs. 9.9±2.4%, n=7 mice, paired t-test, p=0.028, Fig. 2d). The change in magnitude did not correspond to altered reliability between early and late phases, as measured by the coefficient of variation across trials (12.4±1.4 vs. 14.3±0.8, n=7 mice, paired t-test, p=0.239, Fig. 2e). To identify which behavioral components drove the reduced responsiveness, we carried out control experiments with separate cohorts receiving either (1) daily imaging without visual stimulation or training, (2) full training without imaging, or (3) two weeks of visual stimulus exposure without imaging or association with the air-puff. Comparison of visually evoked responses on Day 0 versus Day 15 revealed a significant difference for groups 2 and 3 (Supplemental Figure 2a), indicating that visual experience alone is sufficient to induce a reduction in sensory-evoked cortical activity. To determine the stage of visual processing in which this functional plasticity occurred, we imaged the activity of thalamocortical axons within layer 4 arising from the lateral geniculate nucleus before and after behavioral training (see Supplemental Table 1, Supplemental Figure 2b-c). These results showed a small but non-significant increase in thalamic output, arguing that the experience-dependent reduction in V1 responses arises through modification of intracortical circuits.

We next investigated the ability of a linear classifier to predict the presence of a visual stimulus versus baseline spontaneous activity for individual trials (see Online Methods). Using a support vector machine (SVM) model, we found that individual PNs exhibited a range of accuracy levels that did not significantly differ across learning (early vs. late phase, 0.59±0.02 vs. 0.56±0.01, n=7 mice, paired t-test, p=0.999, Kolmogorov-Smirnoff test, p=0.956, Fig. 2f), despite the substantial reduction in average response magnitude. Interestingly, the stimulus-predictive accuracy of single neurons was significantly correlated with response magnitude within a single learning phase (early: Spearman’s r^2^=0.52, p<0.001; mid: Spearman’s r^2^=0.50, p<0.001; late: Spearman’s r^2^=0.46, p<0.001, Fig. 2G), a somewhat surprising result given the overall reduction in magnitude without loss of accuracy across training. Previous work has shown that neuronal populations can perform much better than individual cells in predicting sensory stimuli ^30^. Therefore, we trained a similar SVM model using an ensemble vector comprising all neurons. Our results confirmed that, as a group, layer 2/3 PNs performed significantly better than chance during all learning phases (early: 0.90±0.03, n=7 mice, t-test, p<0.001; mid: 0.84±0.04, n=7 mice, t-test, p<0.001; late: 0.87±0.03, n=7 mice, t-test, p<0.001, Fig. 2h). In addition, population accuracy was unchanged over learning despite the reduced response magnitude (early vs. late, n=7 mice, paired t-test, p=0.772, Fig. 2h). Overall, these findings demonstrate that evoked activity in V1 is strongly shaped by visual experience, but such functional plasticity can occur without significant alteration in the ability to robustly encode sensory input.

Next, we examined whether visually evoked activity in V1 was predictive of an animal’s behavioral performance. During early and mid learning, there was no difference between the average PN response magnitude for correct versus incorrect trials (early: 21.9±6.3% vs. 20.7±6.9%, n=7 mice, paired t-test, p=0.188; mid: 20.1±7.2% vs. 16.2±7.8%, n=7 mice, paired t-test, p=0.235). However, during the late phase, responses on correct trials were significantly larger than on incorrect (10.8±2.7% vs. 7.8±2.7%, n=7 mice, paired t-test, p=0.003, Fig. 3a-b). To determine if neuronal activity could also predict individual trial outcomes, we again used a linear model to classify responses. Interestingly, the average accuracy of individual neurons increased modestly but significantly over the 2-week training period (early vs. late phase, 0.51±0.004 vs. 0.52±0.003, n=7 mice, paired t-test, p=0.039, Kolmogorov-Smirnoff test, p<0.001, Fig. 3c). In contrast to stimulus prediction accuracy, behavioral accuracy was not correlated with response magnitude for any learning phase (early: Spearman’s r^2^<0.001, p=0.228; mid: Spearman’s r^2^=0.007, P=0.082; late: Spearman’s r^2^=0.020, p=0.086, Fig. 3d). The layer 2/3 PN ensemble was also not better than chance during the early phase of training (0.52±0.01, n=7 mice, t-test, p=0.053), yet did perform better than chance for later phases (mid: 0.57±0.01, n=7 mice, t-test, p<0.001; late: 0.57±0.01, n=7 mice, t-test, p<0.001, Fig. 3e). Moreover, the blink prediction accuracy was significantly better for the late versus early phase (paired t-test, p=0.017).

**Figure 3.**
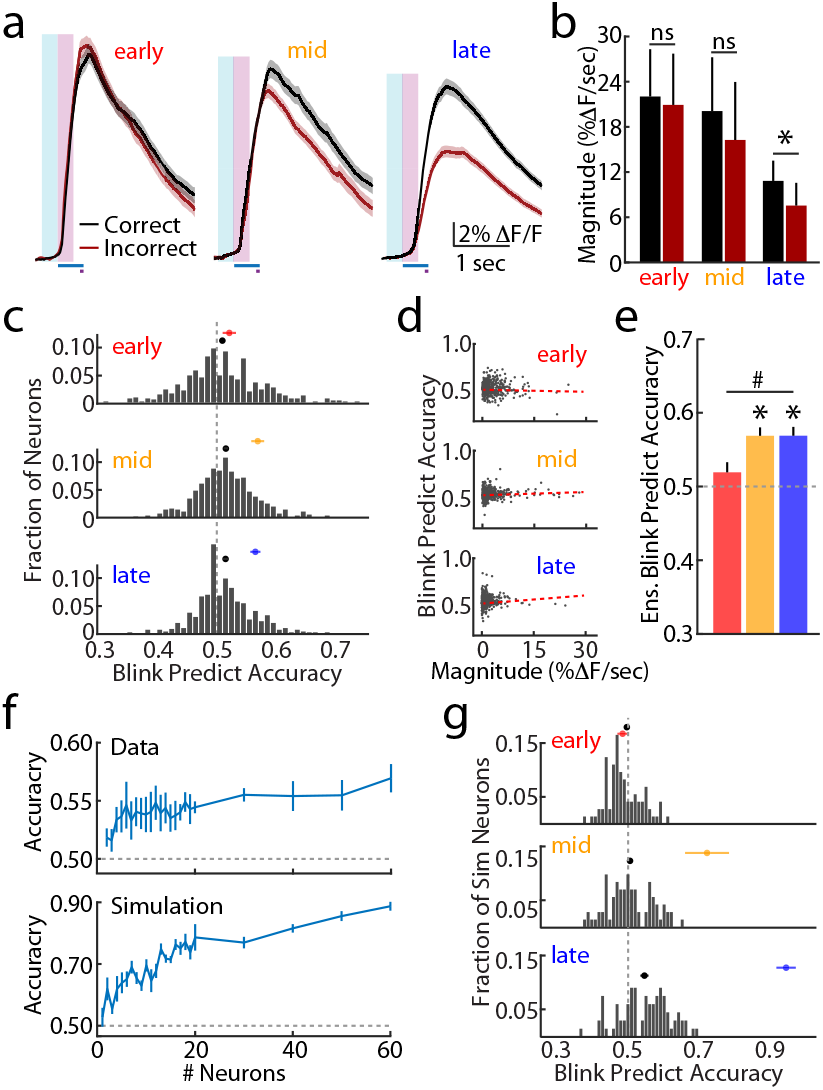
Cortical representation of behavioral outcome emerges during training. **a**, Average visually evoked response for one example layer 2/3 neuron across training phases. Traces are separated by correct (black) and incorrect (dark red) trials. Timing of visual stimulus (blue bar) and air-puff (purple bar), and analysis windows (baseline, light blue; visual response, pink) are shown. Lines and shading indicate average ± SEM across all trials for the given phase. **b**, Average population values for the visual response magnitude, separated by correct and incorrect trials, within each training phase. Bars represent average ± SEM (n=7 mice). * indicates p<0.05 and ns indicates p>0.05, paired t-test for correct vs. incorrect. **c**, Distribution of blink prediction accuracy values using a linear decoder for responses of individual layer 2/3 neurons across each training phase. Chance level (0.5) is indicated (gray dashed line). Black circles indicate average ± SEM for the population of individual neurons. **d**, Relationship between blink prediction accuracy and response magnitude for individual neurons across each training phase. Red dashed line indicates Spearman’s rank correlation. **e**, Average blink prediction accuracy values using a linear decoder for the ensemble activity. Colors denote training phases as above. Bars represent average ± SEM (n=7 mice) and are also indicated by colored circles in (c). * indicates p<0.05, t-test relative to chance for each phase. # indicates p<0.05, paired t-test for early vs. late. **f**, Blink prediction accuracy versus number of cells included in the ensemble for a linear decoder. Chance level (0.5) is indicated (gray dashed line). Upper panel is for data from layer 2/3 neurons. Lower panel is for simulated values with similar group statistics. **g**, Distribution of blink prediction accuracy values using a linear decoder for simulated responses of individual neurons, with group statistics similar to actual data, across each training phase. Chance level (0.5) is indicated (gray dashed line). Black circles indicate average ± SEM for the population of simulated neurons. Colored circles indicate average ± SEM performance of the linear decoder using the ensemble of simulated neurons.

Our results showed that the ensemble could better predict behavioral performance in comparison to the average accuracy of individual neurons. We therefore estimated the number of single cells necessary to reach population-level accuracy by training our classifier on randomly selected neuronal groups of varying size. This analysis suggested that ~30 cells were sufficient to match the accuracy of the overall population (Fig. 3f). We then asked whether the enhanced performance of the ensemble could be explained simply by the pooling of multiple single cells. We simulated a population of neurons whose accuracy values matched the distribution of the layer 2/3 PNs and trained our classifier on these data. As with real neurons, the ensemble showed markedly better performance than the average single cells and a corresponding improvement with learning phase (early vs. late, 0.47±0.01 vs. 0.95±0.02, n=80 neurons, t-test, p<0.001, Fig. 3g). Furthermore, ensemble accuracy levels were reached by pooling ~30 simulated cells (Fig. 3f). Thus, our results demonstrate the surprising finding that the accurate representation of behavior in V1 is highly plastic, emerging during learning despite the unchanged accuracy of stimulus prediction. Moreover, improved ensemble accuracy emerges from the increasing accuracy of individual neurons through pooling of a relatively small number of cells.

Thus far, our data provide evidence for substantial functional plasticity of activity in V1 layer 2/3 neurons during behavioral training, including changes in response magnitude and behavioral encoding. As the output of cortical PNs is strongly shaped by GABAergic interneurons, we investigated how similar training affected local inhibitory circuits in V1. We used AAV vectors to conditionally express GCaMP6f in either PV-, SOM-, or VIP-INs (see Online Methods) and carried out similar behavioral conditioning and imaging as described above (Fig. 4a).

**Figure 4.**
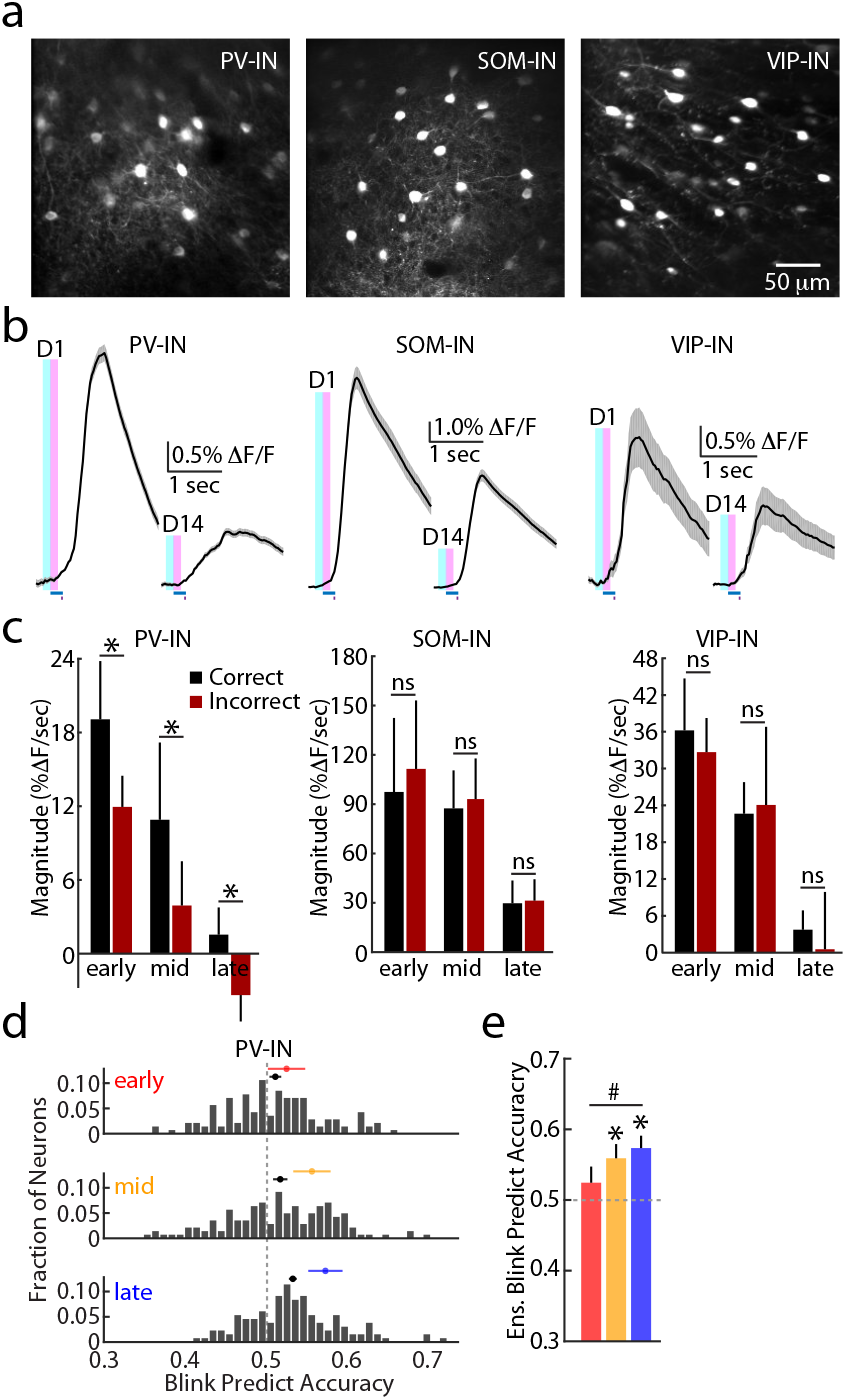
Sensory and behavioral representation by GABAergic interneurons. **a**, Example in vivo 2-photon imaging of GCaMP6f in layer 2/3 PV-INs (left), SOM-INs (middle), and VIP-INs (right). **b**, Average visually evoked responses for example layer 2/3 PV-INs (left), SOM-INs (middle), and VIP-INs (right). Traces are shown for Day 1 and Day 14. Timing of visual stimulus (blue bar) and air-puff (purple bar), and analysis windows (baseline, light blue; visual response, pink) are shown. Lines and shading indicate average ± SEM across all trials. **c**, Average population values for the visual response magnitude, separated by correct (black) and incorrect (dark red) trials, within each training phase. Data are for PV-INs (left), SOM-INs (middle), and VIP-INs (right). Bars represent average ± SEM (PV-INs, n=8 mice; SOM-INs, n=7 mice; VIP-INs, n=7 mice). * indicates p<0.05, paired t-test for correct vs. incorrect. **d**, Distribution of blink prediction accuracy values using a linear decoder for responses of individual layer 2/3 PV-INs across each training phase. Chance level (0.5) is indicated (gray dashed line). Black circles indicate average ± SEM for the population of individual neurons. **e**, Average blink prediction accuracy values using a linear decoder for the ensemble activity of PV-INs. Bars represent average ± SEM (n=8 mice) and are also indicated by colored circles in (d). * indicates p<0.05, t-test relative to chance for each phase. # indicates p>0.05 and p<0.05, respectively, paired t-test for early vs. late.

Task acquisition did not differ across the various strains of mice (Supplemental Figure 3a). Analysis of visually evoked responses revealed that all three interneuron groups exhibited strong and significant reduction in response magnitude over the course of training, similar to results from PNs (Fig. 4b, Supplemental Fig. 3b). As above, this change did not alter the ability of PV-, SOM-, or VIP-IN ensembles to significantly predict the visual stimulus in each training phase (Supplemental Fig. 3c). However, only the PV-INs demonstrated the emergence of a significant difference between response magnitude on correct versus incorrect trials (early: 20.1±4.5% vs. 12.6±2.4%, n=8 mice, paired t-test, p=0.031; mid: 11.1±6.6% vs. 4.2±3.3%, n=8 mice, paired t-test, p=0.047; late: 1.2±2.1% vs. −3.9±2.1%, n=8 mice, paired t-test, p=0.021, Fig. 4c). Moreover, the linear classifier revealed that individual PV-INs exhibited a range of blink prediction accuracy values that significantly increased with training (early vs. late, 0.51±0.01 vs. 0.53±0.004, n=8 mice, paired t-test, p=0.013, Kolmogorov-Smirnoff test, p=0.004, Fig. 4d).

Training the linear classifier on interneuron ensemble data showed that blink prediction accuracy for PV-INs did not differ from chance during early training (0.52±0.02, n=8 mice, t-test, p=0.172) but did perform significantly above chance for mid (0.56±0.02, n=8 mice, t-test, p=0.023) and late (0.57±0.02, n=8 mice, t-test, p=0.009, Fig. 4e). Moreover, there was a significant increase in accuracy between early and late training (paired t-test, p=0.039). Similar analyses for SOM- and VIP-INs revealed that behavioral prediction accuracy did not differ from chance at any phase of training (Supplemental Fig. 3d).

The functional plasticity of V1 activity associated with behavioral training might arise from modification of synaptic connectivity within local networks. NMDARs are strongly linked to synaptic plasticity of both excitatory and inhibitory synapses and may be necessary for experience-dependent changes in visually evoked responses ^10, 31–34^. To determine whether our results are dependent on NMDAR signaling, we used an AAV vector to delete the obligatory GluN1 subunit from a sparse number of V1 neurons. We previously demonstrated the efficacy of this approach for inducing loss of functional NMDARs ^31^. Here, expression of Cre recombinase-tdTomato in GluN1f/f mice allowed us to identify putative GluN1-null cells during in vivo imaging (Fig. 5a-b). On average, 20.8±3.0 of cells in each field of view were identified as Cre-positive.

**Figure 5.**
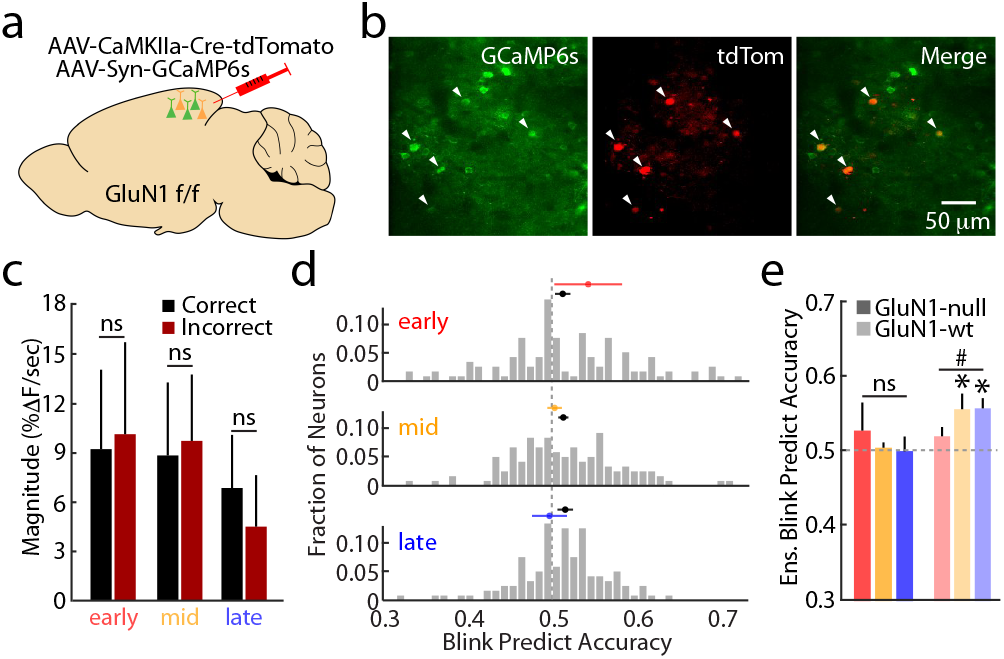
NMDARs are required for functional plasticity of cortical representations. **a**. Schematic illustration showing viral strategy for sparse deletion of the GluN1 subunit and expression of GCaMP6s. **b**, Example in vivo 2-photon image of GCaMP6s (green, left), tdTomato (red, middle), and merge (right) in layer 2/3 neurons. tdTomato-expressing cells are putative GluN1-null. **c**, Average population values for the visual response magnitude of GluN1-null cells, separated by correct (black) and incorrect (dark red) trials, within each training phase. Bars represent average ± SEM (n=6 mice). ns indicates p>0.05, paired t-test for correct vs. incorrect. **d**, Distribution of blink prediction accuracy values using a linear decoder for responses of individual GluN1-null cells across each training phase. Chance level (0.5) is indicated (gray dashed line). Black circles indicate average ± SEM for the population of individual neurons. **e**, Average blink prediction accuracy values using a linear decoder for the ensemble activity of GluN1-null cells (dark bars) and GluN1-wt cells (light bars). Bars represent average ± SEM (n=6 mice) and are also indicated by colored circles in (f). * indicates p<0.05, t-test relative to chance for each phase. ns and # indicate p>0.05 and p<0.05, respectively for early vs. late.

Importantly, sparse loss of NMDARs did not disrupt acquisition of conditioned behavior (Supplemental Fig. 4a). Imaging identified GluN1-null cells revealed no significant reduction in either spontaneous or visually evoked activity across training (see Supplemental Table 1, Supplemental Fig. 4b-d). Training the linear classifier on GluN1-null ensemble data showed that were capable of predicting a visual stimulus above chance for all phases, and this accuracy did not change during training (Supplemental Fig. 4e). Strikingly, GluN1-null cells failed to develop a significant difference in response magnitude for correct versus incorrect trials (early: 9.3±5.1% vs. 10.2±5.4%, n=6 mice, paired t-test, p=0.703; mid: 9.0±4.2% vs. 9.9±4.2%, n=6 mice, paired t-test, p=0.686; late: 6.6±3.3% vs. 4.5±3.0%, n=6 mice, paired t-test, p=0.073, Fig. 5c). Training the linear classifier on data from GluN1-null cells revealed that the average blink prediction accuracy of single neurons did not improve with learning (early vs. late, 0.51±0.01 vs. 0.52±0.01, n=6 mice, paired t-test, p=0.386, Kolmogorov-Smirnoff test, p=0.155, Fig. 5d). Additionally, the GluN1-null ensemble could not predict behavior above chance for any phase (early: 0.52±0.04, n=6 mice, t-test, p=0.294; mid: 0.50±0.01, n=6 mice, t-test, p=0.362; late: 0.50±0.02, n=6 mice, t-test, p=0.557), and accuracy did not improve over training (early vs. late, paired t-test, p=0.694, Fig. 5e).

Given the small number of GluN1-null cells imaged per animal, it may be that prediction accuracy was low simply due to insufficient size of the analysis pool. To examine this possibility, we trained our classifier on a population of 20 randomly selected GluN1-wt neurons (tdTomato-negative) from the same mice as GluN1-null cells. These results revealed that this smaller pool of wild-type cells performed similarly to the overall control population. The GluN1-wt neurons could not predict behavior during the early phase (0.52±0.01, n=7 mice, t-test, p=0.055) but performed significantly better than chance for mid (0.55±0.02, n=7 mice, t-test, p=0.029) and late (0.56±0.01, n=7 mice, t-test, p=0.001) phases and demonstrated significant improvement with training (early vs. late, paired t-test, p=0.026, Fig. 5e). Thus, the inability of GluN1-null neurons to significantly represent behavioral outcome is not due to their smaller overall number, but likely reflects loss of underlying cellular mechanisms linked to functional reorganization of network activity.

## Discussion

Here, we showed that sensory-evoked neuronal activity in V1 is highly plastic during visual experience associated with acquisition of a conditioned eyeblink task. Response magnitude for both excitatory and inhibitory cells was strongly reduced over several days. However, this decrease occurred without loss of accuracy for predicting the sensory stimulus. In marked contrast, the accurate representation of behavioral outcome (correct versus incorrect) on individual trials emerged gradually during learning for both PNs and PV-INs. These results are consistent with previous work indicating that sensory experience, both passive or associated with task learning, can drive plasticity in cortical networks and alter the representation of sensory information ^5–8, 11, 12^. In most cases, such studies have shown an enhancement of tuning specificity, suggesting that experience can lead to sharpened selectivity for behaviorally relevant stimuli. In contrast, our data show that classical conditioning can selectively modify the representation of behavioral output. Earlier studies from our lab and others demonstrate the ability of stimulus-evoked neuronal activity in primary sensory cortex to accurately encode an animal’s subsequent behavioral choice ^6, 21, 22, 35^, though whether this representation is present initially or only emerges during experience was unclear. Interestingly, visual learning can drive the experience-dependent expression of reward signals in V1, further supporting this form of behaviorally-relevant functional plasticity ^19^. Overall, these results highlight the convergence of information streams within V1, corresponding to feed-forward sensory input, top-down signals of behavioral state, and neuromodulatory influences that all interact to dynamically modify output early in the sensory processing hierarchy.

Our finding that visual experience, independent of training, drives a strong suppression of sensory response magnitude potentially contrasts with earlier work showing that visually evoked potentials (VEPs) that largely reflect thalamocortical synaptic input can exhibit enhanced amplitude in response to repeated presentation of patterned stimuli ^11^. However, our results from imaging thalamic axons showed a similar, though not statistically significant, enhancement in evoked activity associated with training. Thus, our findings support a model in which increased thalamic input occurs alongside suppression of layer 2/3 activity. The mechanisms for these opposing results are unclear. Increased output from GABAergic interneurons might account for intracortical suppression, though this hypothesis is inconsistent with our observation that all three major classes of inhibitory cells exhibit similar decreased response magnitude. Another likely candidate mechanism is experience-dependent modification of synaptic strength. We show that sparse deletion of the obligatory GluN1 subunit of the NMDAR blocks both the suppression of visually evoked responses and the emergence of accurate behavioral representation. As NMDAR signaling is linked to plasticity of both excitatory and inhibitory connections ^32, 33^, this finding supports the hypothesis that synaptic changes contribute to functional plasticity in vivo.

Although GluN1 deletion abolished both modification of response magnitude and behavioral representation, our results suggest these processes may be independent, as response magnitude was not correlated with prediction accuracy. Future investigations may allow us to fully disentangle the cellular and circuit mechanisms underlying both forms of functional plasticity. Importantly, the finding that NMDARs appear to act cell-autonomously (neighboring GluN1-wt cells exhibit normal behavioral representation) indicates that the experience-dependent difference between visually evoked response magnitude on correct and incorrect trials does not simply reflect a change in the downstream correlation of V1 activity and motor output. Instead, intracortical plasticity may be a fundamental contributor to the learning process.

We observed emergence of accurate behavioral predictions in both PNs and PV-INs, but not in SOM- or VIP-INs, likely reflecting cell type-specificity of underlying plasticity mechanisms. PV-IN activity is closely linked to that of local excitatory networks and plays a key role in regulating the timing and gain of sensory-evoked responses ^36–38^. Thus, the functional coupling of these populations is not surprising. Interestingly, SOM-INs are linked to the control of dendritic calcium signaling and experience-dependent circuit plasticity ^5, 39–41^. However, we find that these cells do not exhibit accurate behavioral predictions. Nevertheless, SOM- and VIP-IN output may be central to the mechanisms underlying modification of neuronal activity, a hypothesis that awaits further experimental testing.

In conclusion, our results strongly support a role for primary sensory cortex in visually guided behavior. Earlier studies showed that cortical circuits are required for normal task performance ^21, 42, 43^, though other work suggests minimal contribution of the neocortex to perceptual ability ^44^. The present data indicate that at a minimum, V1 networks can accurately represent behavior, enabling the transmission of such information to multiple downstream areas engaged during both learning and expert performance. Moreover, the dynamic modification of neuronal output and prediction accuracy during training highlights the critical role of experience in shaping behaviorally relevant activity in the neocortex.

## Acknowledgements

The authors are extremely thankful to Dr. Lan Tang for initial design of the eyeblink conditioning task and helpful comments during analyses. We also wish to thank Dr. Jessica A. Cardin as well as Mr. Daniel Barson, Dr. Tom Morse, and Dr. Garrett Neske, and other members of the Higley Laboratory for helpful comments during the preparation of this manuscript.

## Authors contributions

Experiments were conceived and designed by AP and MJH. Data were acquired by AP. Analyses and interpretation were designed and carried out by AP, HB, and MJH. Manuscript was written by AP and MJH.

## Data and materials availability

All data are presented in the paper and supplementary materials. The datasets generated during the current study are available from the corresponding author on reasonable request.

## Online Methods

### Animals

Animals were handled in accordance with the Yale Institutional Animal Care and Use Committee and federal guidelines. C57BL/6 mice were purchased from Envigo. *PV*^Cre^/ C57/BL6 (Jackson laboratory, RRID: IMSR_JAX:008069), *SST*^Cre^/ C57/BL6 (Jackson laboratory, RRID: IMSR_JAX:013044), *VIP*^Cre^/ C57/BL6 (Jackson laboratory, RRID: IMSR_JAX:031628) and GluN1^fl/fl^ (Jackson laboratory, RRID: IMSR_JAX: 005246) mice were bred in-house from animals originally purchased from Jackson Laboratory. Animals of both sexes were used and aged 8-10 weeks old at the beginning of the experimental procedures. All mice were group housed (2-3 same-sex animals per cage) under a 12 h/12 h light/dark cycle with water and food provided *ad libitum*. From the day of the first stereotaxic surgery, animals were fed sulfatrim mouse chow (Uniprim). All experiments were performed during the light phase of the daily cycle. In all housing and experimental rooms, the temperature was maintained at 23-24°C, with humidity levels between 35% and 45%.

### Surgery

Five weeks prior to imaging and behavioral experiments, mice were injected stereotactically with an adenoassociated viral (AAV) vector driving expression of a genetically encoded calcium indicator either in non-specific neuron populations (AAV5-syn-GCamP6s, C57BL/6 mice), targeted populations of interneurons (AAV5-syn-flex-GCamp6f, *PV*^Cre^/ C57/BL6, *SST*^Cre^/ C57/BL6 or *VIP*^Cre^/ C57/BL6 mice), or thalamocortical axonal terminals arising from the lateral geniculate nucleus (AAV5-syn-jGCaMP7b, C57BL/6 mice). To examine the effects of deletion of the GluN1 subunit of the NMDA-type glutamate receptor, GluN1^fl/fl^ animals were injected with AAV5-syn-GCamP6s and dilute AAV5-EF1a-iCre-TdTomato (Baylor Vector Core, 1:300 in saline). Mice were anesthetized with isoflurane and received subcutaneous injection of an analgesic and anti-inflammatory drug (Carprofen 2mg/ml in saline, 5ml/kg). Mice were then placed in a stereotaxic apparatus (David Kopf Instruments) and their scalp shaved and disinfected with 70% ethanol. Ocular lubricant was used to protect animals’ eyes from drying during surgery. To deliver the viruses into the left visual cortex (V1, coordinates AP: −0.35 cm; LM: −0.25 cm; DV: −0.055 cm) and dorsal lateral geniculate nucleus (dLGN, coordinates AP: −0.235 cm; LM: −0.2 cm; DV: −0.29 cm), we used a Nanofil 36G beveled needle inserted through a small craniotomy. The syringe was connected to a Micro Syringe Pump (World Precision Instruments) used to deliver virus (0.5-0.7 μl or 0.2 μl of total volume for cortical or thalamic injections respectively, 100 nl/min). After the injection, the needle remained in the brain for 5 min to allow for diffusion of the virus. Seven to ten days after viral injections, animals were implanted with cranial windows and titanium head-posts. Subjects were anesthetized with isoflurane and received subcutaneous injection of an analgesic and anti-inflammatory drug (Carprofen 2mg/ml in saline, 5ml/kg). Skin and periosteum were reflected and the skull was cleaned with saline and dried. Two screws were set into the skull over the right hemisphere, and a custom-made titanium headpost (~2g) was fixed to the bone with dental cement (Metabond, Parkell). A craniotomy (approx. 4 mm^2^) was made over the left V1 and a bilayer cranial window (5×5mm No. 1 cover glass and 3.5×3.5 No. 1 cover glass, bonded using ultraviolet-curing adhesive, Norland Products) was inserted into the opening and fixed to the skull using instant glue (Krazy Glue) and dental cement (Metabond, Parkell).

### Behavioral Setup

The mouse was head-fixed on a freely-moving wheel (15 cm diameter) under the objective of a 2-photon microscope located in a light-proof chamber. Visual stimuli were displayed on a computer monitor positioned normal to and 22 cm away from the right eye. Air-puffs (10-12 psi) were delivered to the right cornea via a small metal cannula coupled to a compressed air tank and gated by a solenoid (Clark Solutions). Timing of the air puff was coordinated with the visual stimulus using custom-written MATLAB codes through a NI-DAQmx board (PCIe-6315, National Instruments) at a sampling rate of 5 kHz. Eyelid closure and pupil diameter were continuously recorded using a monochromatic CMOS camera (PointGrey FlyCapture3) at a frame rate of 33 fps. An infrared LED array was used to illuminate the eye. All signals, including the timing of the visual stimuli, the air puffs, the wheel position, video frame ticks, and microscope resonant scanner frame ticks were digitized (5 kHz) and collected through a Power 1401 (CED) acquisition board using Spike 2 software.

### Behavioral Training

Starting nine days prior to training, mice were habituated to head-fixation while placed on a freely-moving running wheel (15 cm diameter), gradually increasing from a few minutes to one hour over this period. After habituation, training consisted of 75 daily presentations to the right eye of a 500 ms visual stimulus (CS+) presented on a gamma-corrected monitor (20° sinusoidally drifting grating, 0.05 cycles per degree, 1 cycle per second, 100% contrast). For each animal, the stimulus location was fixed in one of nine 3×3 sub-regions of the screen that evoked the largest population response in the field of view. Each stimulus co-terminated with a 50 ms air-puff directed to the ipsilateral cornea. Training was carried out over 14 consecutive days (Days 1-14). In addition to this protocol, on the day preceding training (Day 0) and the day following training (Day 15), each animal was presented with 50 CS presentations in the absence of a coupled air-puff. These three trial types were randomly interleaved. For all training days, the inter-trial interval was 9-13 seconds, with each trial value randomly selected from a flat hazard distribution.

### Calcium Imaging

Imaging was carried out using a two-photon Movable Objective Microscope (MOM) with a galvo-resonant scanner (Sutter Instruments) through a 25x, 1.05 NA objective (Olympus) coupled to a Ti-sapphire laser (MaiTai eHP DeepSee, SpectraPysics) tuned to 920 nm. Collection of tdTomato images were carried out at 1000 nm. Images were acquired using ScanImage 2017 (Vidrio) at ~30 Hz and a resolution of 256×256. The microscope and the perimeter of the objective were tightly wrapped in blackout material to prevent light contamination from the LCD screen. Somata of layer 2/3 neurons were imaged at approximately 180-300 μm depth relative to the brain surface^45^. Thalamocortical axons in layer 4 were imaged at 330-460 μm depth.

### Behavioral Analysis

Eyeblink videos were analyzed with custom MATLAB scripts as previously described ^21^. Briefly, gray-scale images from each training session were binarized to maximize the contrast between the eye (white) and surrounding fur (black). A region of interest around the eye was manually defined, and the time-varying proportion of white to dark pixels was used as a readout of eye closure. These data were normalized by the 5th and 95th percentile values for each session, resulting in a range of 0 to 1, corresponding to a fully open and fully closed eye, respectively. The conditioned response (CR) was defined as the maximum eye closure during the 450 ms window between visual stimulus and air-puff onset. The unconditioned response (UR) was defined as the maximum eye closure within a 500 ms window from the onset of the air-puff. Trials were identified as correct if the CR:UR ratio was larger than 0.1. Trials were excluded from analysis if the eye closed >10% within a 2-second window prior to visual stimulus onset. Spontaneous blinks (>10% eye closure) were detected during the inter-trial-intervals. The spontaneous blink rate was calculated as the average number of blinks per 450 ms interval to compare with the behavioral analysis window.

### Imaging Analysis

Images of neuronal activity were first motion-corrected using the Moco plugin for ImageJ ^46^. The first 150 frames of each movie were used as a template with the maximal distance to be translated in the x and y directions between 20 and 40 pixels. Videos from successive days were translated onto the first-day template. Regions of interest (ROIs) were selected as previously described ^47^. Further data analysis was performed using custom MATLAB scripts. Fluorescence (F) over time was measured by averaging within the ROI, and contamination from the surrounding neuropil was removed with a discounting coefficient of 0.7^45, 47^. ΔF/F was calculated as (F-F_0_)/F_0_, where F_0_ was the lowest 10% of values from the neuropil-subtracted trace for each session.

To relate neuronal activity to behavior, we divided the data into 3 distinct learning phases of equal duration based on average performance: early (days 1 to 3), mid (days 4 to 6), and late (days 12-14). All data were grouped within a single phase. The magnitude of the visual response on each trial was defined in one of two ways: (1) the mean ΔF/F in a 300ms time window after visual stimulus onset, subtracting the mean ΔF/F over the 300 ms preceding the stimulus or (2) the slope of the visual response measured as the linear fit to ΔF/F within 300ms window following visual stimulus onset. Machine learning was used to decode neuronal activity and assess the accuracy of predicting either the visual stimulus or the conditioned response. A linear Support Vector Machine (SVM) classifier was trained and tested by using an available Matlab toolkit, libsvm, with 5-fold cross-validation and bootstrapping to achieve balanced labels for correct versus incorrect trials. To determine prediction accuracy for the visual stimulus, for each trial we quantified the slope of ΔF/F for the 300ms preceding and following the visual stimulus onset. We also identified a similar matched pair of values obtained for a randomly selected pseudo-onset during the inter-stimulus period. The model was trained two classify the presence of a visual stimulus (i.e., distinguish trials from pseudo-trials). A similar approach was used to determine prediction accuracy for single cells and for the population. To determine prediction accuracy for the conditioned response, we used only paired slope values corresponding to actual stimulus onset times and trained the model to classify correct versus incorrect trials. Again, this approach was used to determine single cell and population performance. To determine how population size influences behavioral prediction accuracy, we repeatedly trained the model on randomly drawn neuronal subsets of varying size. To investigate the relationship between average single neuron performance and population performance, we simulated a distribution of 80 independent slope values for 75 trials, matching the means and variances of the actual neuronal data on correct and incorrect trials and across learning phases. We then used these simulated values to train the same SVM model and assess simulated accuracy.

### Statistics

For all statistical comparisons, values were averaged within each animal, and final analyses were performed with animal number as the degree of freedom. We opted for this approach given the inherent lack of independence for cells imaged within the same animal. The analysis of simulated neuronal performance was an exception to this approach, given the inherent structure of the data. Statistical tests included paired and one-sample T-tests using an alpha value of 0.05, Kolmogorov-Smirnoff tests, and Spearman’s rank correlation.

**Supplemental Figure 1.**
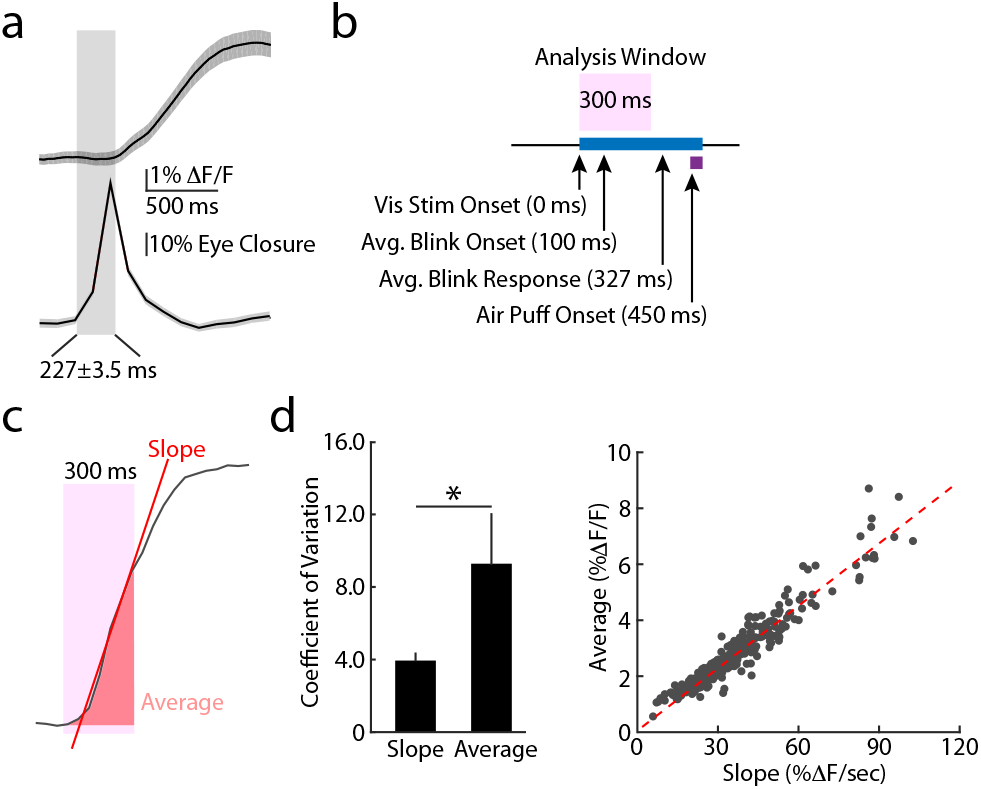
Analysis of visually evoked responses. **a**, Average eyelid trajectory (lower trace) and associated calcium transient (upper trace) corresponding to spontaneous blinks. Gray bar highlights the average latency from blink onset to calcium response. **b**, Schematic illustrating the analysis windows used for all measures of visual responses. The pink bar highlights the window following visual stimulus onset and before contamination of any blink- or air-puff-associated signal. **c**, Average population values for the coefficient of variation, measured for either response slope or average during the 300 ms analysis window for all neurons. Bars represent average ± SEM. * indicates p<0.05, paired t-test. **d**, Relationship between the mean average and slope measures of visual response for each neuron. Red dashed line indicates Spearman’s rank correlation.

**Supplemental Figure 2.**
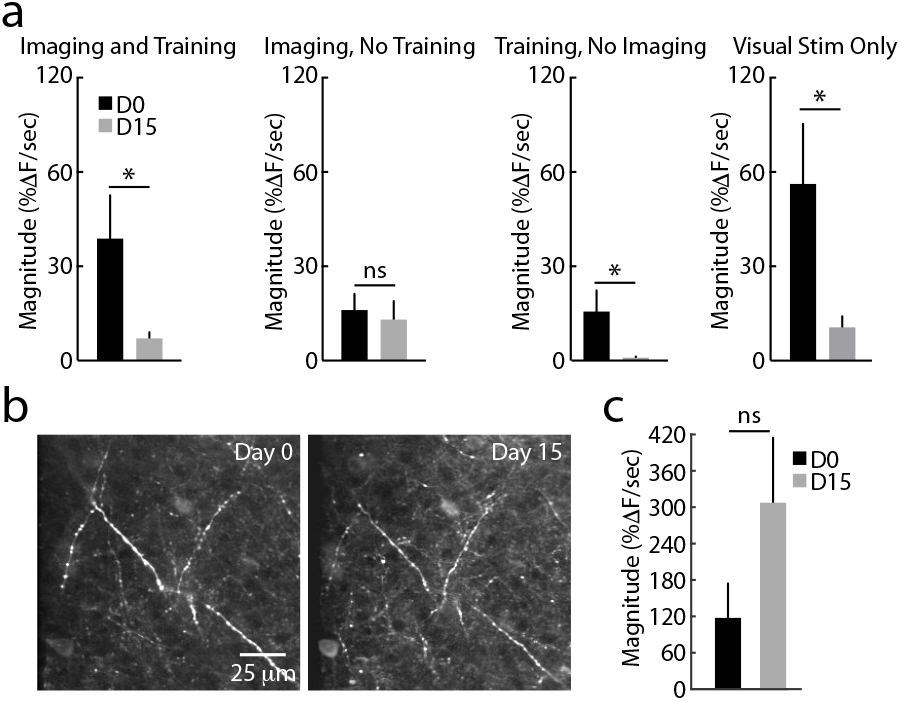
Visual experience drives intracortical plasticity of sensory-evoked responses. **a**, Average population values for the visual response magnitude of layer 2/3 neurons, measured on Day 0 and Day 15. Data are for animals experiencing normal training and imaging (1^st^ panel, replicated from Figure. 2), daily imaging without behavioral training (2^nd^ panel), normal training without imaging (3^rd^ panel), and visual stimulation without air-puff or imaging (4^th^ panel). Bars represent average ± SEM. ns and * indicate p>0.05 or p<0.05, respectively, paired t-test. **b**, Example in vivo 2-photon image of GCaMP6s-expressing thalamocortical axons in layer 4, collected on Day 0 and Day 15. **c**, Average population values for the visual response magnitude of thalamocortical axons, measured on Day 0 and Day 15 from animals receiving normal training. Bars indicate average ± SEM. ns indicates p>0.05, paired t-test.

**Supplemental Figure 3.**
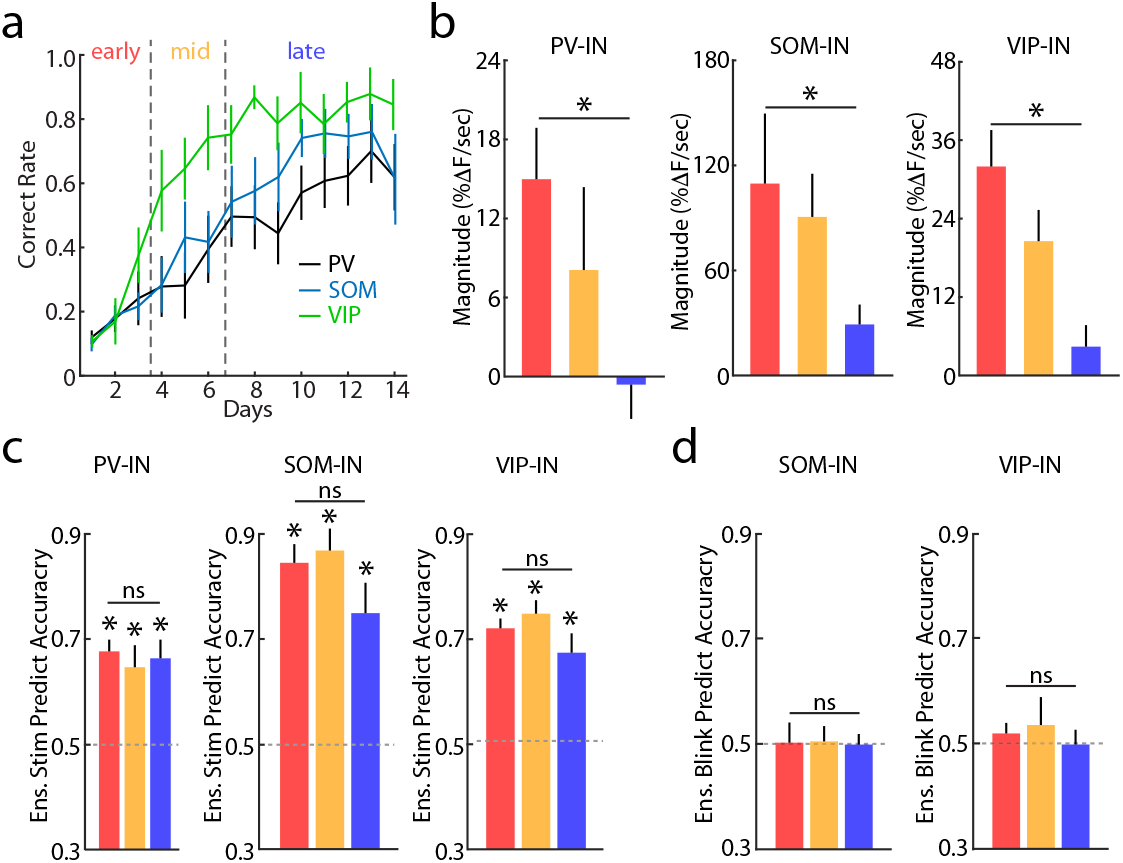
Behavior and associated neuronal responses in GABAergic interneurons. **a**, Behavioral performance over learning for three interneuron imaging cohorts (PV-INs, n=7 mice, black; SOM-INs, n=7 mice, blue; VIP-INs, n=7 mice, green). Lines with error bars represent average ± SEM. **b**, Average population values for response magnitude within each training phase, corresponding to colors in (a). Data are for PV-INs (left), SOM-INs (middle), and VIP-INs (right). Bars represent average ± SEM (n=7 mice for each group). * indicates p<0.05, paired t-test for early vs. late. **c**, Average stimulus prediction accuracy values using a linear decoder for the ensemble activity of PV-INs (left), SOM-INs (middle), and VIP-INs (right). Bars represent average ± SEM (n=8 mice for PV-INs, n=7 mice for SOM- and PV-INs). * indicates p<0.05, t-test relative to chance for each phase. ns indicates p>0.05, paired t-test for early vs. late. **d**, Average blink prediction accuracy values using a linear decoder for the ensemble activity of SOM-INs (left) and VIP-INs (right). Bars represent average ± SEM (n=7 mice for each group). ns indicates p>0.05, paired t-test for early vs. late.

**Supplemental Figure 4.**
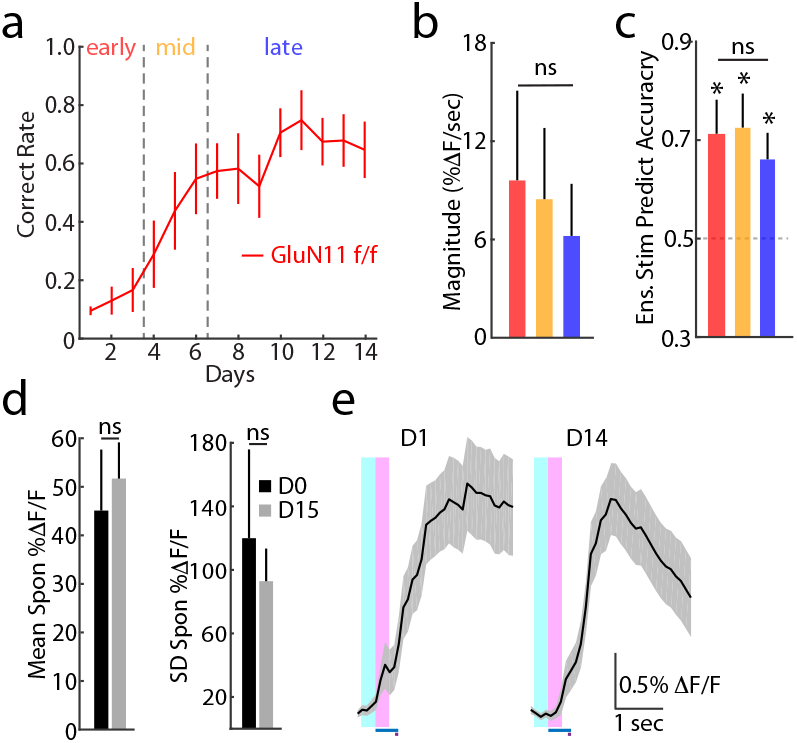
Behavior and associated neuronal responses for GluN1-null cells. **a**, Behavioral performance over learning for mice with sparse deletion of GluN1 in V1 (n=6 mice). Lines with error bars represent average ± SEM. **b**, Average population values for response magnitude within each training phase, corresponding to colors in (a). Bars represent average ± SEM (n=6 mice). ns indicates p>0.05, paired t-test for early vs. late. **c**, Average stimulus prediction accuracy values using a linear decoder for the ensemble activity of GluN1-null cells. Bars represent average ± SEM (n=6 mice). * indicates p<0.05, t-test relative to chance for each phase. ns indicates p>0.05, paired t-test for early vs. late. **d**, Mean (left) and standard deviation (right) of spontaneous activity (%ΔF/F) averaged across all GluN1-null cells, for Day 1 (black) and Day 15 (gray). Bars represent average ± SEM. ns indicates p>0.05, paired t-test. **e**, Average visually evoked responses for example layer 2/3 GluN1-null neuron. Traces are shown for Day 1 (left) and Day 14 (right). Timing of visual stimulus (blue bar) and air-puff (purple bar), and analysis windows (baseline, light blue; visual response, pink) are shown. Lines and shading indicate average ± SEM across all trials.

**Supplemental Table 1.** Summary of all statistical analyses.

| Figure | Comparison | Test | Statistic A | Value A | Statistic B | Value B | Test statistic | 95% Confidence Interval | p-value |
| --- | --- | --- | --- | --- | --- | --- | --- | --- | --- |
| **Imaging** |  |  |  |  |  |  |  |  |  |
| Fig.1f | Spontaneous activity (%ΔF/F), Day 1 vs. Day 14 (n=7 mice) | Paired t-test | Mean %ΔF/F, Day 1 | 44.3+/-5.1% | Mean %ΔF/F, Day 14 | 41.7+/-7.9% |  |  | 0.797 |
| Fig.1f | Standard deviation of spontaneous activity (%ΔF/F), Day 1 vs. Day 14 (n=7 mice) | Paired t-test | Standard deviation %ΔF/F, Day 1 | 70.5+/-25.2% | Standard deviation %ΔF/F, Day 14 | 61.2+/-20.6% |  |  | 0.817 |
| Fig.S1d | CV of slope (%ΔF/sec) vs. CV of average (%ΔF/F) (n=7 mice) | Paired t-test | CV of slope, Day 0 | 4.0 +/- 0.4 | CV of average, Day 0 | 9.29+/-2.62 |  |  | 0.044 |
| Fig.S1d | Correlation of mean (%ΔF/F) and slope (%ΔF/sec) (n=643 cells) | Spearman's rank correlation | Mean (%ΔF/F), Day 0 | n/a | Mag (%ΔF/sec), Day 0 | n/a | r<sup>2</sup>=0.87 |  | <0.001 |
| Fig.2d | Magnitude (%ΔF/sec) for early vs. late phase of training (n=7 mice) | Paired t-test | Mag (%ΔF/sec), early | 21.0+/-6.9% | Mag (%ΔF/sec), late | 9.9+/-2.4% |  |  | 0.028 |
| Fig.2e | CV of magnitude (%ΔF/sec) in early vs. late phase of training (n=7 mice) | Paired t-test | CV of mag (%ΔF/sec), early | 12.4+/-1.4 | CV of slope (%ΔF/sec), late | 14.3+/-0.8 |  |  | 0.239 |
| Fig.2f | Single cell accuracy for stimulus prediction on early vs. late phase of training (n=7 mice) | Paired t-test | Single cell accuracy for stimulus prediction, early | 0.59+/-0.02 | Single cell accuracy for stimulus prediction, late | 0.56+/-0.01 |  |  | 0.999 |
| Fig.2f | Distribution of single cell accuracy for stimulus prediction on early vs. late phase of training (n=7 mice) | KS test | Distribution of single cell accuracy for stimulus prediction, early | n/a | Distribution of single cell accuracy for stimulus prediction, late | n/a |  |  | 0.956 |
| Fig.2g | Single cell accuracy for stimulus prediction vs. magnitude (%ΔF/sec) in early phase of training (n=643 cells) | Spearman's rank correlation | Single cell accuracy for stimulus prediction, early | n/a | Mag (%ΔF/sec), early | n/a | r<sup>2</sup>=0.52 |  | <0.001 |
| Fig.2g | Single cell accuracy for stimulus prediction vs. magnitude (%ΔF/sec) in mid phase of training (n=643 cells) | Spearman's rank correlation | Single cell accuracy for stimulus prediction, mid | n/a | Mag (%ΔF/sec), mid | n/a | r<sup>2</sup>=0.50 |  | <0.001 |
| Fig.2g | Single cell accuracy for stimulus prediction vs. magnitude (%ΔF/sec) in late phase of training (n=643 cells) | Spearman's rank correlation | Single cell accuracy for stimulus prediction, late | n/a | Mag (%ΔF/sec), late | n/a | r<sup>2</sup>=0.46 |  | <0.001 |
| Fig.2h | Ensemble accuracy for stimulus prediction on early vs. late phase of training (n=7 mice) | Paired t-test | Ensemble accuracy for stimulus prediction, early | 0.90+/-0.03 | Ensemble accuracy for stimulus prediction, late | 0.865+/-0.030 |  |  | 0.772 |
| Fig.2h | Ensemble accuracy for stimulus prediction on early, mid and late phase of training vs. chance (n=7 mice) | t-test | Ensemble accuracy for stimulus prediction, early | 0.90+/-0.03 | Chance level | 0.50 |  |  | <0.001 |
|  |  | t-test | Ensemble accuracy for stimulus prediction, mid | 0.84+/-0.04 | Chance level | 0.50 |  |  | <0.001 |
|  |  | t-test | Ensemble accuracy for stimulus prediction, late | 0.87+/-0.03 | Chance level | 0.50 |  |  | <0.001 |
| Fig.S2a | Magnitude (%ΔF/sec) on Day 0 vs. Day 15, Group imaging and training (n=7 mice) | Paired t-test | Imaging and training mag (%ΔF/sec), Day 0 | 38.7+/-13.8% | Imaging and training mag (%ΔF/sec), Day 15 | 7.2+/-1.5% |  |  | 0.026 |
| Fig.S2a | Magnitude (%ΔF/sec) on Day 0 vs. Day 15, Group imaging, no training (n=3 mice) | Paired t-test | Imaging, no training mag (%ΔF/sec), Day 0 | 24.3+/-7.8% | Imaging, no training mag (%ΔF/sec), Day 15 | 19.8+/-9.0% |  |  | 0.096 |
| Fig.S2a | Magnitude (%ΔF/sec) on Day 0 vs. Day 15, Group training, no imaging (n=3 mice) | Paired t-test | Training, no imaging mag (%ΔF/sec), Day 0 | 24.0+/-10.5% | Training, no imaging mag (%ΔF/sec), Day 15 | 4.5+/-0.6% |  |  | 0.002 |
| Fig.S2a | Magnitude (%ΔF/sec) on Day 0 vs. Day 15, Group visual stimulus only (n=3 mice) | Paired t-test | Visual stimulus only mag (%ΔF/sec), Day 0 | 56.4+/-19.2% | Visual stimulus only mag (%ΔF/sec), Day 15 | 10.5+/-3.6% |  |  | 0.030 |
| Fig.S2c | Axon magnitude (%ΔF/sec) on Day 0 vs. Day 15 (n=3 mice) | Paired t-test | Axon mag (%ΔF/sec), Day 0 | 117.1%+/-57.3% | Axon mag (%ΔF/sec), Day 15 | 306.4+/-108.0% |  |  | 0.085 |
| Fig.3b | Magnitude (%ΔF/sec) on correct vs. incorrect trials in early phase of training (n=7 mice) | Paired t-test | Mag (%ΔF/sec) on correct trials, early | 21.9+/-6.3% | Mag (%ΔF/sec) on incorrect trials, early | 20.7+/-6.9% |  |  | 0.188 |
| Fig.3b | Magnitude (%ΔF/sec) on correct vs. incorrect trials in mid phase of training (n=7 mice) | Paired t-test | Mag (%ΔF/sec) on correct trials, mid | 20.1+/-7.2% | Mag (%ΔF/sec) on incorrect trials, mid | 16.2+/-7.8% |  |  | 0.235 |
| Fig.3b | Magnitude (%ΔF/sec) on correct vs. incorrect trials in late phase of training (n=7 mice) | Paired t-test | Mag (%ΔF/sec) on correct trials, late | 10.8+/-2.7% | Mag (%ΔF/sec) on incorrect trials, late | 7.8+/-2.7% |  |  | 0.003 |
| Fig.3c | Single cell accuracy for blink prediction on early vs. late phase of training (n=7 mice) | Paired t-test | Single cell accuracy for blink prediction, early | 0.51+/-0.004 | Single cell accuracy for stimulus prediction, late | 0.52+/-0.003 |  |  | 0.039 |
| Fig.3c | Distribution of single cell accuracy for blink prediction on early vs. late phase of training (n=7 mice) | KS test | Distribution of single cell accuracy for blink prediction, early | n/a | Distribution of single cell accuracy for blink prediction, late | n/a |  |  | <0.001 |
| Fig.3d | Single cell accuracy for blink prediction vs. magnitude (%ΔF/sec) in early phase of training (n=643 cells) | Spearman's rank correlation | Single cell accuracy for stimulus prediction, early | n/a | Mag (%ΔF/sec), early | n/a | r<sup>2</sup><0.001 |  | 0.228 |
| Fig.3d | Single cell accuracy for blink prediction vs. magnitude (%ΔF/sec) in mid phase of training (n=643 cells) | Spearman's rank correlation | Single cell accuracy for stimulus prediction, mid | n/a | Mag (%ΔF/sec), mid | n/a | r<sup>2</sup>=0.007 |  | 0.082 |
| Fig.3d | Single cell accuracy for blink prediction vs. magnitude (%ΔF/sec) in late phase of training (n=643 cells) | Spearman's rank correlation | Single cell accuracy for stimulus prediction, late | n/a | Mag (%ΔF/sec), late | n/a | r<sup>2</sup>=0.020 |  | 0.086 |
| Fig.3e | Ensemble accuracy for blink prediction on early vs. late phase of training (n=7 mice) | Paired t-test | Ensemble accuracy for blink prediction, early | 0.52+/-0.01 | Ensemble accuracy for stimulus prediction, late | 0.57+/-0.01 |  |  | 0.017 |
| Fig.3e | Ensemble accuracy for blink prediction on early, mid and late phase of training vs. chance (n=7 mice) | T-test | Ensemble accuracy for blink prediction, early | 0.52+/-0.01 | Chance level | 0.500 |  |  | 0.053 |
|  |  | T-test | Ensemble accuracy for blink prediction, mid | 0.57+/-0.01 | Chance level | 0.500 |  |  | <0.001 |
|  |  | T-test | Ensemble accuracy for blink prediction, late | 0.57+/-0.01 | Chance level | 0.500 |  |  | <0.001 |
| Fig.3g | Simulated ensemble accuracy for blink prediction on early vs. late phase of training (n=7 mice) | Paired t-test | Simulated ensemble accuracy for blink prediction, early | 0.47+/-0.01 | Simulated ensemble accuracy for blink prediction, late | 0.95+/-0.02 |  |  | <0.001 |
| Fig.3g | Simulated single cell accuracy for blink prediction on early vs. late phase of training (n=7 mice) | Paired t-test | Simulated single cell accuracy for blink prediction, early | 0.49+/-0.01 | Simulated single cell accuracy for blink prediction, late | 0.55+/-0.01 |  |  | <0.001 |
| Fig.3g | Distribution of simulated single cell accuracy for blink prediction on early vs. late phase of training (n=7 mice) | KS test | Distribution of simulated single cell accuracy for blink prediction, early | n/a | Distribution of simulated single cell accuracy for blink prediction, late | n/a |  |  | <0.001 |
| Fig.S3b | PV-IN magnitude (%ΔF/sec) for early vs. late phase of training (n=8 mice) | Paired t-test | PV-IN mag (%ΔF/sec), early | 15.0+/-3.9% | PV-IN mag (%ΔF/sec), late | -0.6+/-2.4% |  |  | 0.003 |
| Fig.S3b | SOM-IN magnitude (%ΔF/sec) for early vs. late phase of training (n=7 mice) | Paired t-test | SOM-IN mag (%ΔF/sec), early | 108.4+/-42.3% | SOM-IN mag (%ΔF/sec), late | 30.0+/-12.4% |  |  | 0.032 |
| Fig.S3b | VIP-IN magnitude (%ΔF/sec) for early vs. late phase of training (n=7 mice) | Paired t-test | VIP-IN mag (%ΔF/sec), early | 31.5+/-5.4% | VIP-IN mag (%ΔF/sec), late | 4.2+/-3.3% |  |  | 0.026 |
| Fig.S3c | PV-IN ensemble accuracy for stimulus prediction on early vs. late phase of training (n=8 mice) | Paired t-test | PV-IN ensemble accuracy for stimulus prediction, early | 0.69+/-0.04 | PV-IN ensemble accuracy for stimulus prediction, late | 0.65+/-0.03 |  |  | 0.810 |
| Fig.S3c | PV-IN ensemble accuracy for stimulus prediction on early, mid and late phase of training vs. chance (n=8 mice) | T-test | PV-IN ensemble accuracy for stimulus prediction, early | 0.69+/-0.04 | Chance level | 0.500 |  |  | 0.002 |
|  |  | T-test | PV-IN ensemble accuracy for stimulus prediction, mid | 0.64+/-0.03 | Chance level | 0.500 |  |  | 0.006 |
|  |  | T-test | PV-IN ensemble accuracy for stimulus prediction, late | 0.65+/-0.03 | Chance level | 0.500 |  |  | 0.003 |
| Fig.S3c | SOM-IN ensemble accuracy for stimulus prediction on early vs. late phase of training (n=7 mice) | Paired t-test | SOM-IN ensemble accuracy for stimulus prediction, early | 0.83+/-0.06 | SOM-IN ensemble accuracy for stimulus prediction, late | 0.77+/-0.01 |  |  | 0.902 |
| Fig.S3c | SOM-IN ensemble accuracy for stimulus prediction on early, mid and late phase of training vs. chance (n=7 mice) | T-test | SOM-IN ensemble accuracy for stimulus prediction, early | 0.83+/-0.06 | Chance level | 0.500 |  |  | <0.001 |
|  |  | T-test | SOM-IN ensemble accuracy for stimulus prediction, mid | 0.85+/-0.08 | Chance level | 0.500 |  |  | <0.001 |
|  |  | T-test | SOM-IN ensemble accuracy for stimulus prediction, late | 0.77+/-0.01 | Chance level | 0.500 |  |  | 0.006 |
| Fig.S3c | VIP-IN ensemble accuracy for stimulus prediction on early vs. late phase of training (n=7 mice) | Paired t-test | VIP-IN ensemble accuracy for stimulus prediction, early | 0.72+/-0.02 | VIP-IN ensemble accuracy for stimulus prediction, late | 0.67+/-0.04 |  |  | 0.897 |
| Fig.S3c | VIP-IN ensemble accuracy for stimulus prediction on early, mid and late phase of training vs. chance (n=7 mice) | T-test | VIP-IN ensemble accuracy for stimulus prediction, early | 0.72+/-0.02 | Chance level | 0.500 |  |  | <0.001 |
|  |  | T-test | VIP-IN ensemble accuracy for stimulus prediction, mid | 0.75+/-0.03 | Chance level | 0.500 |  |  | <0.001 |
|  |  | T-test | VIP-IN ensemble accuracy for stimulus prediction, late | 0.67+/-0.04 | Chance level | 0.500 |  |  | 0.003 |
| Fig.S3d | SOM-IN ensemble accuracy for blink prediction on early vs. late phase of training (n=7 mice) | Paired t-test | SOM-IN ensemble accuracy for blink prediction, early | 0.50+/-0.04 | SOM-IN ensemble accuracy for blink prediction, early | 0.50+/-0.02 |  |  | 0.527 |
| Fig.S3d | SOM-IN ensemble accuracy for blink prediction on early, mid and late phase of training vs. chance (n=7 mice) | T-test | SOM-IN ensemble accuracy for blink prediction, early | 0.50+/-0.04 | Chance level | 0.500 |  |  | 0.488 |
|  |  | T-test | SOM-IN ensemble accuracy for blink prediction, mid | 0.50+/-0.02 | Chance level | 0.500 |  |  | 0.451 |
|  |  | T-test | SOM-IN ensemble accuracy for blink prediction, late | 0.50+/-0.02 | Chance level | 0.500 |  |  | 0.543 |
| Fig.S3d | VIP-IN ensemble accuracy for blink prediction on early vs. late phase of training (n=7 mice) | Paired t-test | VIP-IN ensemble accuracy for blink prediction, early | 0.52+/-0.03 | SOM-IN ensemble accuracy for blink prediction, late | 0.50+/-0.03 |  |  | 0.887 |
| Fig.S3d | VIP-IN ensemble accuracy for blink prediction on early, mid and late phase of training vs. chance (n=7 mice) | T-test | VIP-IN ensemble accuracy for blink prediction, early | 0.52+/-0.03 | Chance level | 0.500 |  |  | 0.264 |
|  |  | T-test | VIP-IN ensemble accuracy for blink prediction, mid | 0.54+/-0.05 | Chance level | 0.500 |  |  | 0.264 |
|  |  | T-test | VIP-IN ensemble accuracy for blink prediction, late | 0.50+/-0.03 | Chance level | 0.500 |  |  | 0.560 |
| Fig.4c | PV-IN magnitude (%ΔF/sec) on correct vs. incorrect trials in early phase of training (n=8 mice) | Paired t-test | PV-IN mag (%ΔF/sec) on correct trials, early | 20.1+/-4.5% | PV-IN mag (%ΔF/sec) on incorrect trials, early | 12.6+/-2.4% |  |  | 0.031 |
| Fig.4c | PV-IN magnitude (%ΔF/sec) on correct vs. incorrect trials in mid phase of training (n=8 mice) | Paired t-test | PV-IN mag (%ΔF/sec) on correct trials, mid | 11.1+/-6.6% | PV-IN mag (%ΔF/sec) on incorrect trials, mid | 4.2+/-3.3% |  |  | 0.047 |
| Fig.4c | PV-IN magnitude (%ΔF/sec) on correct vs. incorrect trials in late phase of training (n=8 mice) | Paired t-test | PV-IN mag (%ΔF/sec) on correct trials, late | 1.2+/-2.1% | PV-IN mag (%ΔF/sec) on incorrect trials, late | -3.9+/-2.1% |  |  | 0.021 |
| Fig.4c | SOM-IN magnitude (%ΔF/sec) on correct vs. incorrect trials in early phase of training (n=7 mice) | Paired t-test | SOM-IN mag (%ΔF/sec) on correct trials, early | 99.1+/-45.2% | SOM-IN mag (%ΔF/sec) on incorrect trials, early | 108.4+/-42.3% |  |  | 0.983 |
| Fig.4c | SOM-IN magnitude (%ΔF/sec) on correct vs. incorrect trials in mid phase of training (n=7 mice) | Paired t-test | SOM-IN mag (%ΔF/sec) on correct trials, mid | 87.3+/-24.1% | SOM-IN mag (%ΔF/sec) on incorrect trials, mid | 93.2+/-27.2% |  |  | 0.931 |
| Fig.4c | SOM-IN magnitude (%ΔF/sec) on correct vs. incorrect trials in late phase of training (n=7 mice) | Paired t-test | SOM-IN mag (%ΔF/sec) on correct trials, late | 30.1+/-12.2% | SOM-IN mag (%ΔF/sec) on incorrect trials, late | 33.3+/-12.4% |  |  | 0.828 |
| Fig.4c | VIP-IN magnitude (%ΔF/sec) on correct vs. incorrect trials in early phase of training (n=7 mice) | Paired t-test | VIP-IN mag (%ΔF/sec) on correct trials, early | 36.1+/-9.2% | VIP-IN mag (%ΔF/sec) on incorrect trials, early | 33.4+/-6.1% |  |  | 0.320 |
| Fig.4c | VIP-IN magnitude (%ΔF/sec) on correct vs. incorrect trials in mid phase of training (n=7 mice) | Paired t-test | VIP-IN mag (%ΔF/sec) on correct trials, mid | 24.1+/-6.4% | VIP-IN mag (%ΔF/sec) on incorrect trials, mid | 24.1+/-12.0% |  |  | 0.546 |
| Fig.4c | VIP-IN magnitude (%ΔF/sec) on correct vs. incorrect trials in late phase of training (n=7 mice) | Paired t-test | VIP-IN mag (%ΔF/sec) on correct trials, late | 3.0+/-3.1% | VIP-IN mag (%ΔF/sec) on incorrect trials, late | -0.3+/-9.1% |  |  | 0.367 |
| Fig.4d | PV-IN single cell accuracy for blink prediction on early vs. late phase of training (n=8 mice) | Paired t-test | PV-IN single cell accuracy for blink prediction, early | 0.51+/-0.01 | PV-IN single cell accuracy for blink prediction, late | 0.53+/-0.004 |  |  | 0.013 |
| Fig.4d | Distribution of PV-IN single cell accuracy for blink prediction on early vs. late phase of training (n=8 mice) | KS test | Distribution of PV-IN single cell accuracy for blink prediction, early | n/a | Distribution of PV-IN single cell accuracy for blink prediction, late | n/a |  |  | 0.004 |
| Fig.4e | PV-IN ensemble accuracy for blink prediction on early vs. late phase of training (n=8 mice) | Paired t-test | PV-IN ensemble accuracy for blink prediction, early | 0.52+/-0.02 | PV-IN ensemble accuracy for blink prediction, late | 0.57+/-0.02 |  |  | 0.039 |
| Fig.4e | PV-IN ensemble accuracy for blink prediction on early, mid and late phase of training vs. chance (n=8 mice) | T-test | PV-IN ensemble accuracy for blink prediction, early | 0.52+/-0.02 | Chance level | 0.500 |  |  | 0.172 |
|  |  | T-test | PV-IN ensemble accuracy for blink prediction, mid | 0.56+/-0.02 | Chance level | 0.500 |  |  | 0.023 |
|  |  | T-test | PV-IN ensemble accuracy for blink prediction, late | 0.57+/-0.02 | Chance level | 0.500 |  |  | 0.009 |
| Fig.S4b | GluN1-null magnitude (%ΔF/sec) for early vs. late phase of training (n=6 mice) | Paired t-test | GluN1-null mag (%ΔF/sec), early | 9.6+/-5.4% | GluN1-null mag (%ΔF/sec), late | 6.3+/-3.3% |  |  | 0.090 |
| Fig.S4c | GluN1-null ensemble accuracy for stimulus prediction on early, mid and late phase of training vs. chance (n=6 mice) | Paired t-test | GluN1-null ensemble accuracy for stimulus prediction, early | 0.71+/-0.07 | GluN1-null ensemble accuracy for stimulus prediction, late | 0.65+/-0.06 |  |  | 0.789 |
| Fig.S4c | GluN1-null ensemble accuracy for stimulus prediction on early, mid and late phase of training vs. chance (n=6 mice) | T-test | GluN1-null ensemble accuracy for stimulus prediction, early | 0.71+/-0.07 | Chance level | 0.500 |  |  | 0.012 |
|  |  | T-test | GluN1-null ensemble accuracy for stimulus prediction, mid | 0.73+/-0.07 | Chance level | 0.500 |  |  | 0.013 |
|  |  | T-test | GluN1-null ensemble accuracy for stimulus prediction, late | 0.65+/-0.06 | Chance level | 0.500 |  |  | 0.029 |
| Fig.S4d | GluN1-null spontaneous activity (%ΔF/F), Day 1 vs. Day 14 (n=6 mice) | Paired t-test | GluN1-null mean %ΔF/F, Day 1 | 45.1+/-1.3% | GluN1-null mean %ΔF/F, Day 14 | 51.7+/-7.5% |  |  | 0.723 |
| Fig.S4d | GluN1-null standard deviation of spontaneous activity (%ΔF/F), Day 1 vs. Day 14 (n=6 mice) | Paired t-test | GluN1-null standard deviation %ΔF/F, Day 1 | 119.9+/-5.6% | GluN1-null standard deviation %ΔF/F, Day 14 | 92.9+/-2.1% |  |  | 0.692 |
| Fig.5c | GluN1-null magnitude (%ΔF/sec) on correct vs. incorrect trials in early phase of training (n=6 mice) | Paired t-test | GluN1-null mag (%ΔF/sec) on correct trials, early | 9.3+/-5.1% | GluN1-null mag (%ΔF/sec) on incorrect trials, early | 10.2+/-5.4% |  |  | 0.703 |
| Fig.5c | GluN1-null magnitude (%ΔF/sec) on correct vs. incorrect trials in mid phase of training (n=6 mice) | Paired t-test | GluN1-null mag (%ΔF/sec) on correct trials, mid | 9.0+/-4.2% | GluN1-null mag (%ΔF/sec) on incorrect trials, mid | 9.9+/-4.2% |  |  | 0.686 |
| Fig.5c | GluN1-null magnitude (%ΔF/sec) on correct vs. incorrect trials in late phase of training (n=6 mice) | Paired t-test | GluN1-null mag (%ΔF/sec) on correct trials, late | 6.6+/-3.3% | GluN1-null mag (%ΔF/sec) on incorrect trials, late | 4.5+/-3.0% |  |  | 0.073 |
| Fig.5d | GluN1-null single cell accuracy for blink prediction on early vs. late phase of training (n=6 mice) | Paired t-test | GluN1-null single cell accuracy for blink prediction, early | 0.51+/-0.01 | GluN1-null single cell accuracy for blink prediction, late | 0.52+/-0.01 |  |  | 0.386 |
| Fig.5d | Distribution of GluN1-null single cell accuracy for blink prediction on early vs. late phase of training (n=6 mice) | KS test | Distribution of GluN1-null single cell accuracy for blink prediction, early | n/a | Distribution of GluN1-null single cell accuracy for blink prediction, late | n/a |  |  | 0.155 |
| Fig.5e | GluN1-null ensemble accuracy for blink prediction on early, mid and late phase of training vs. chance (n=6 mice) | Paired t-test | GluN1-null ensemble accuracy for blink prediction, early | 0.52+/-0.04 | GluN1-null ensemble accuracy for blink prediction, late | 0.50+/-0.02 |  |  | 0.694 |
| Fig.5e | GluN1-wt ensemble (20 cells) accuracy for blink prediction on early, mid and late phase of training vs. chance (n=6 mice) | Paired t-test | GluN1-wt ensemble (20 cells) accuracy for blink prediction, early | 0.52+/-0.01 | GluN1-wt ensemble (20 cells) accuracy for blink prediction, late | 0.56+/-0.01 |  |  | 0.026 |
| Fig.5e | GluN1-null ensemble accuracy for blink prediction on early, mid and late phase of training vs. chance (n=6 mice) | T-test | GluN1-null ensemble accuracy for blink prediction, early | 0.52+/-0.04 | Chance level | 0.500 |  |  | 0.294 |
|  |  | T-test | GluN1-null ensemble accuracy for blink prediction, mid | 0.50+/-0.01 | Chance level | 0.500 |  |  | 0.362 |
|  |  | T-test | GluN1-null ensemble accuracy for blink prediction, late | 0.50+/-0.02 | Chance level | 0.500 |  |  | 0.557 |
| Fig.5e | GluN1-wt ensemble (20 cells) accuracy for blink prediction on early, mid and late phase of training vs. chance (n=6 mice) | T-test | GluN1-wt ensemble (20 cells) accuracy for blink prediction, early | 0.52+/-0.01 | Chance level | 0.500 |  |  | 0.055 |
|  |  | T-test | GluN1-wt ensemble (20 cells) accuracy for blink prediction, mid | 0.55+/-0.02 | Chance level | 0.500 |  |  | 0.029 |
|  |  | T-test | GluN1-wt ensemble (20 cells) accuracy for blink prediction, late | 0.56+/-0.01 | Chance level | 0.500 |  |  | 0.001 |

